# Compressive Stress Enhances Invasive Phenotype of Cancer Cells via Piezo1 Activation

**DOI:** 10.1101/513218

**Authors:** Mingzhi Luo, Kenneth K. Y. Ho, Zhaowen Tong, Linhong Deng, Allen P. Liu

## Abstract

Uncontrolled growth in solid tumor generates compressive stress that drives cancer cells into invasive phenotypes, but little is known about how such stress affects the invasion and matrix degradation of cancer cells and the underlying mechanisms. Here we show that compressive stress enhanced invasion, matrix degradation, and invadopodia formation of breast cancer cells. We further identified Piezo1 channels as the putative mechanosensitive cellular components that transmit the compression to induce calcium influx, which in turn triggers activation of RhoA, Src, FAK, and ERK signaling, as well as MMP-9 expression. Interestingly, for the first time we observed invadopodia with matrix degradation ability on the apical side of the cells, similar to those commonly observed at the cell’s ventral side. Furthermore, we demonstrate that Piezo1 and caveolae were both involved in mediating the compressive stress-induced cancer cell invasive phenotype as Piezo1 and caveolae were often colocalized, and reduction of Cav-1 expression or disruption of caveolae with methyl-β-cyclodextrin led to not only reduced Piezo1 expression but also attenuation of the invasive phenotypes promoted by compressive stress. Taken together, our data indicate that mechanical compressive stress activates Piezo1 channels to mediate enhanced cancer cell invasion and matrix degradation that may be a critical mechanotransduction pathway during, and potentially a novel therapeutic target for, breast cancer metastasis

## Introduction

Cancer invasion is a cumulative result of multiple processes including directed cell migration and extracellular matrix (ECM) degradation. While chemical factors are well known in mediating cancer invasion, mechanical forces such as compressive stress have also been identified as essential regulators of these processes^1^. For example, an increase of compressive stress inside solid tumor accompanies with cell proliferation and stiffness enhancement^2–5^. Cancer cells also experience compressive stress when they migrate through capillary and confined tissue microenvironment^6–9^. Reports have shown that compressive stress alters cell-cell attachment, cell adhesion, traction force, proliferation, differentiation, and migration^2,4,10,11^. Recent *in vivo* studies show that compressive stress stimulates tumorigenic signaling by increasing β-catenin signaling in colon epithelial cells^12^, and strategies to release compressive stress can indeed enhance the efficiency of anti-tumor treatment^13^. Interestingly, it is demonstrated *in vitro* that compressive stress directly drives cancer cells to invasive phenotypes by forming leading cells^14^. However, whether compressive stress enhances invasive phenotypes by promoting both migration and matrix degradation of the cancer cells and how the compressive stress is sensed and transduced into cellular behaviors is still poorly understood.

Considering that compressive stress stretches cell membrane and thus increases membrane tension, it may as well alter cancer cells’ behaviors through tension-mediated conformational changes of proteins and lipids in the membrane^15^. In particular, the increase of membrane tension can activate several stretch-activated ion channels (SAC) including Piezo and transient receptor potential (TRP) channels^16–18^. Comparing to other SAC such as TRP channels that are also involved in the sensing and response to mechanical stress^19^, Piezo channels are known to respond to membrane tension with more exquisite sensitivity^20^. In synthetic lipid bilayers, mechanical stress can activate purified Piezo channels even in the absence of other cellular components^21^. This and other work convincingly show that Piezo channels are extremely mechanosensitive^21,22^. On the other hand, studies *in vivo* show that Piezo channels mediate diverse physiological activities that are associated with compressive stimulation including touch perception^18^, blood pressure sensing^23^, vascular development^24^ and breathing^25,26,27^, and are essential in some mechanically related pathological processes such as pressure-induced pancreatitis^28^ and breast cancer development^29^. In the case of breast cancer development, the role of Piezo1 is substantiated by the shorter survival times of patient with upregulated Piezo1 mRNA expression level^29^. More importantly, it has been shown that the breast cancer cells’ response to compression is dependent on Piezo but not TRP channels^30^, and upregulation of Piezo1 mRNA expression leads to a stronger response to negative pressure in breast cancer cells as compared with their normal counterparts^29^. These data are consistent with that compressive stress promotes the migration of cancer cells but not normal cells^14^ and indicate that Piezo1 may be essential for the compression-induced enhancement of cancer invasion. However, whether and how Piezo1 channels mediate compressive stress-enhanced invasive phenotype of cancer cells has not been examined.

So far it is generally known that SAC functions at “membrane force foci” such as caveolae^31^, as the cholesterol- and sphingolipid-enriched caveolae are thought to provide proper platforms for harboring and gating SAC^21,32^. In fact, several ion channels harbored in caveolae of myocytes are known to regulate mechanotransduction during cell swelling^33^. The specific structure of caveolae may provide a situation in which the change of membrane tension is different from the surrounding membrane when cells are exposed to mechanical stress. Indeed, it has been found that increase of membrane tension due to mechanical stress can rapidly flatten and disassemble the flask-like membrane invaginations characteristic of caveolae as a compensation mechanism^34,35^, and TRP channels such as TRPC1 are formed in signal complex to facilitate the interaction with specific lipids such as cholesterol and sphingolipid in caveolae^36^. Structural analysis of Piezo1 has shown that there is a lipid pocket sandwiched between Piezo1 repeat B and C, which provides a lipid binding site and thus may be accessed by lipid molecules as a means of regulation^27^. Despite such evidence of structure for lipid interaction, the localization of Piezo1 is not well established and thus it remains unclear whether its activity is regulated by caveolae.

In this study, we hypothesized that Piezo1 channels mediate the compressive stress-enhanced invasive phenotype of cancer cells *via* a caveolae-dependent mechanism. To test this hypothesis, we examined *in vitro* cultured human breast cancer cells for their ability to migrate and degrade extracellular matrix in the presence or absence of compressive mechanical stress, together with corresponding changes in Piezo1 as well as cytoskeletal remodeling and calcium signaling. We found that compressive stress promoted an invasive phenotype in the breast cancer cells, characterized by enhanced cell migration, invadopodia formation and matrix degradation, stress fiber remodeling and calcium signal initiation. More importantly, the phenotypic changes in these cells appeared to be mediated by compression-induced Piezo1 activation, which in turn was largely dependent on the integrity of caveolae.

## Materials and Methods

### Cell culture and preparation

MDA-MB-231 cells (ATCC HTB-26), an invasive breast adenocarcinoma cell line, were cultured in DMEM with 2 mM L-glutamine (# 11965-092, Thermo Fisher, MA) supplemented with 10% fetal bovine serum (FBS, # 35-010-CV, Thermo Fisher, MA), 100 units/ml penicillin, 100 units/ml streptomycin, 2.5 μg/ml Fungizone, and 5 μg/ml gentamicin (# 15750-060, Invitrogen, MA) at 5% CO_2_ and 37 °C. MCF10A (ATCC^®^ CRL-10317™) cells, a normal breast cell line, were cultured in DMEM/Ham’s F-12 (Gibco-Invitrogen, Carlsbad, CA) supplemented with 100 ng/ml cholera toxin, 20 ng/ml epidermal growth factor (EGF; # PHG0311, Invitrogen, MA), 0.01 mg/ml insulin, 500 ng/ml hydrocortisone, and 10% FBS. For matrix degradation and invadopodia experiments, cells were incubated in invadopodia medium containing DMEM supplement with 5% Nu-Serum (# 355104, Corning, NY), 10% FBS, and 20 ng/ml EGF.

For labeling actin in live cells, stable cell lines expressing Lifeact-RFP were generated *via* lentiviral transfection. The lentiviral transfer plasmids pLVX-puro-GFP-Lifeact and pLVX-puro-RFP-Lifeact were cloned from RFP-Lifeact plasmid obtained from Dr. Gaudenz Danuser (UT-Southwestern). Briefly, lentiviruses were produced by transfecting human embryonic kidney (HEK)-293T cells with psPAX2 and pMD2.G (Addgene) and pLVX-puro-GFP-Lifeact viral vectors. Conditioned medium containing viruses was collected after 5 days and then used immediately to infect cells or stored at −80 °C. Transduced target cells were selected with puromycin for 72h.

For optical imaging of dynamic calcium signaling and caveolae localization in live cells, cell lines transiently expressing G-GECO (a green fluorescent genetically encoded calcium indicator) and Caveolin-1 (Cav-1)-EGFP respectively were generated *via* plasmid transfection. The plasmids expressing G-GECO was a generous gift from Takanari Inoue (Johns Hopkins University)^37^, and those expressing Cav-1-EGFP were from Ari Helenius (ETH Zurich). Briefly, cells were transfected with Lipofectamine-2000 (# 11668-019, Life Technologies). For 35 mm glass bottom dishes, 6 µg plasmid DNA in Optimem^®^ transfection medium (# 31985062, Gibco) was used for each transfection. After 24 h at 37 °C, the transfection medium was replaced with complete medium, and cells were processed after 24-48 h later.

### *In vitro* compression device

To investigate the effect of compressive stress on cell behaviors, we used a previously described setup^14,38^. Briefly, cells were grown either in a 35 mm culture dish with glass bottom (# 12-565-90, Fisher Scientific) that was coated with/without gelatin, or in a transwell chamber with permeable membrane of 8-μm pores that was coated with Matrigel. Then the cells were covered with a soft agarose disk layer, and subsequently a piston of specific weight was placed on top of the agarose disk to apply a given compressive stress to the cells underneath indirectly. The cross-sectional area of the piston (24 mm diameter) was 4.52 cm^2^ but its weight was variable at 9.22 g, 8.45 g, and 27.67 g, corresponding to a stress of 200 Pa, 400 Pa, and 600 Pa, respectively, on the cells. Cells prepared as such but not subjected to piston weight were used as controls.

### *In vitro* transwell invasion assay

To assay the effect of compressive stress on cell invasion, standard transwell invasion assay adapted from Bravo-Cordero^39,40^ was performed using 6-well Transwell chambers that were separated as upper and lower chambers by filter membrane with 8 μm pores (# 07-200-169, Corning). For the assay, the transwell filter membrane was coated with 300 µl Matrigel (12 mg/ml, # E1270, Sigma, Burlington, MA) diluted in serum-free DMEM (2 mg/ml final concentration), followed by incubation for 1 h at 37 °C. MDA-MB-231 cells in serum-free medium (5×10^5^ cells/well) were placed in the upper chamber, while the lower chamber was filled with 2 ml complete medium. Cells were allowed to grow for 6 h and then compressed for 18 h before being fixed with 4% paraformaldehyde (# 30525-89-4, Electron Microscopy Sciences, Hatfield, PA). The non-invasive cells on the upper chamber were removed with cotton swabs, and the invaded cells in the lower chamber were stained with 0.1% crystal violet (# C6158; Sigma) for 10 min at room temperature, prior to being examined and imaged by light microscopy at 10× magnification (Olympus BX60; Olympus Corporation, Tokyo, Japan). Then the number of stained cells in the lower chamber was counted using ImageJ software (National Institute of Health, Bethesda, MD) and the enhancement of cellular invasion induced by compressive stress was quantified as percentage (%) of the number of compressed cells over that of the non-compressed cells that had invaded through the filter membrane, *i.e.* [# of cells in the lower chamber in the presence of a specific weight (experiment group)]/[# of cells in the lower chamber in the absence of a specific weight (control group)]. Results are based on the analysis of 10 random fields per transwell in each condition and each experiment was repeated three times.

### Evaluation of invadopodia formation and ECM degradation

To determine whether compressive stress enhances cells’ ability to degrade ECM, we examined cells cultured on gelatin substrate for their tendency to form invadopodia and associated gelatin degradation, according to a protocol adapted from Artym *et al.*^41^. Briefly, glass bottom dishes were treated with 20% nitric acid for 1 h, washed with H2O for 4 times, then incubated with 50 μg/ml poly-L-lysine (# P8920, Sigma) in PBS for 15 min and washed with PBS, then further incubated with 0.5% glutaraldehyde in PBS on ice for 15 min followed by thorough washes with PBS. Subsequently, the dishes were coated with 1 ml of gelatin in PBS (1: 9 of 0.1% fluorescein isothiocyante (FITC)-gelatin (# G13186, Invitrogen, MA): 2% porcine gelatin), then washed in PBS, incubated with 5 mg/ml sodium borohydride (NaBH4) for 3 min, rinsed in PBS, and then incubated in 10% calf serum/DMEM at 37° for 2 h. Afterwards, cells were seeded in each dish at 5×10^5^ cells per well and incubated for 8 h, and then subjected to compressive stress of either 200 Pa, 400 Pa, or 600 Pa, respectively, for 8 h as aforementioned.

Upon completion of compression, the cells were imaged with live fluorescence microscopy (at 60x) and the microscopic images were analyzed by using ImageJ to assess the formation of invadopodia and the degradation of gelatin matrix. Invadopodia were defined as F-actin-positive puncta protruding from the cells into the gelatin matrix underneath the cell in our experiments^42^. For each independent experiment that was performed in triplicates, the number of invadopodia per cell was quantified with cells imaged randomly in >15 microscope view fields, representing a total of ~100 cells per experimental condition. At the same time, the degradation of gelatin matrix was quantified as the percentage of degraded area (dark spots comprised of dense degraded protein products) in the whole area underneath each cell.

### Live fluorescence microscopy

To observe the dynamics of actin, Cav-1, and calcium signaling, live cells expressing Lifeact-RFP, Cav-1-EGFP, and G-GECO were imaged with a spinning disk confocal microscope with a 100x oil immersion objective (Olympus IX73 with Yokogawa CSU-X1). For live fluorescence microscopy, cells were seeded in a 35 mm glass bottom dish that was placed in an environmental chamber mounted on the microscope to maintain constant 37 °C, 5% CO_2_, and humidity. Cav-1-EGFP was observed at the excitation wavelength of 488 nm. For dynamic tracking of stress fiber and caveolae in live cells, the cells were consecutively imaged for up to 60 min, and the images were processed using ImageJ. Cells were observed from both top-down and side view for spatial localization of invadopodia, stress fibers and caveolae by 3D reconstruction of images in Z-stacks (0.4 μm increments).

### Intracellular Ca^2+^ measurement

To evaluate the intracellular calcium concentration ([Ca^2+^]), we used cells transiently expressed with calcium-sensitive reporter G-GECO^43^ and then evaluated the intensity of intracellular calcium signaling. Briefly, cells transfected with G-GECO for 48 h were plated into a glass bottom dish, which was further incubated for 24 h. Subsequently, the cells were imaged with the spinning disk confocal microscope (60 x objective), with fluorescence excitation and emission at 488 nm and 533 nm, respectively. For each experimental group, twenty cells were randomly selected and the fluorescence intensity per cell was quantified using ImageJ.

### Drug treatments

For experiments involving inhibitors, cells were exposed to inhibitor for 0.5 h, unless stated otherwise, in presence or absence of compressive stress. For inhibiting the function of mechanically sensitive ion channels, cells were treated with Gadolinium chloride (Gd^3+^, 5 μM, # 203289, Sigma, Burlington, MA) or GsMTx4 (5 μM, #ab141871; Abcam, Cambridge, MA). For inhibiting the activity of Src, cells were treated with PP2 (10 μM, Calbiotech, Spring Valley, CA). To remove calcium ion from the DMEM, EGTA (2 mM, # E3889; Sigma, Burlington, MA) was added to the medium. To disrupt caveolae in the membrane, cells were treated with 5 mM of methyl-β-cyclodextrin (MβCD, # SLBP3372V, Sigma, Burlington, MA).

### Antibodies for immunoblotting and Western blot

Antibodies used in immunofluorescence and Western blot include: anti-MMP9 rabbit polyclonal antibody (# AB13458) and anti-GAPDH mouse monoclonal antibody (# CB1001) purchased from EMD Millipore (Billerica, MA); anti-Src rabbit antibody (# 2108), anti-p-Src (Y416) rabbit antibody (# 2101), anti-p44/42 MAPK (ERK1/2) mouse monoclonal antibody (# 4696), and anti-p-ERK1/2 (Thr202Tyr204) rabbit monoclonal antibi (# 4370) obtained from Cell Signaling Technology (Danvers, MA), respectively; anti-cortactin rabbit monoclonal antibody (# Ab81208) purchased from Abcam (Cambridge, MA); Caveolin-1 rabbit polyclonal antibody (# PA1-064) and anti-pFAK (Y397) rabbit monoclonal antibody (# 44624) obtained from Thermo Fisher (Grand Island, NY).

### RNA interference

To silence the expression of Piezo1 and Cav-1, Negative Control Medium GC Duplex #2 and siRNA interference for Piezo1 (# AM16708, Assay ID:138387, Thermo Fisher) and Cav-1 (# AM16708, Assay ID: 10297, Thermo Fisher) were used. Briefly, Cells were seeded in 6-well plates at 1 × 10^6^ cells/well for 24 h prior to transfection. At 90% confluence the cells were transfected with 30 nmol/L siRNA using Lipofectamine RNAi MAX (# 13778, Invitrogen, MA) in OptiMEM according to the manufacturer’s instructions. Transfection mixes were applied to the cells for 24 h, subsequently removed and replaced with 2 ml of growth media. The cells were cultured for 48 h before use in experiments. The protein expression levels of Piezo1 and Cav-1 were ascertained by Western blotting.

### Immunofluorescence

Cells were fixed with 4% paraformaldehyde for 10 min and permeabilized with 0.1 % TritonX-100 for 10 min at room temperature. Non-specific sites were blocked using 5 % non-fat milk in PBS for 1 h at room temperature. Cells were then incubated in 5 % non-fat milk in PBS containing primary antibodies at 1:100 dilution for 1 h at room temperature. After washing with PBS, cells were incubated with Alexa Fluor 594 or 640 conjugated secondary antibody for 60 min at room temperature. Cells were visualized using the spinning disk confocal microscope with 60x oil immersion objective. For F-actin staining, cells were incubated with 1:100 rhodamine phalloidin (# PHDR1, Cytoskeleton Inc.) for 60 min at room temperature.

### G-LISA analysis

To determine the RhoA activity, a G-LISA RhoA Activation Assay Biochem kit (# BK124; Cytoskeleton Inc., Denver, CO, USA) was used according to the manufacturer’s instructions. Briefly, the samples were homogenized in ice-cold lysis buffer (R0278, Sigma) with a protease-inhibitor cocktail, and then centrifuged at 450x g at 4 °C for 1 min. The supernatants were harvested, and protein concentrations were measured using the Precision Red Advanced Protein Assay Reagent (# ADV02; Cytoskeleton Inc.) and finally equalized with ice-cold lysis buffer to 1.0 mg/ml. Equalized protein extractions were transferred to a Rho-GTP-binding protein pre-coated plate (Cytoskeleton, Inc.). The plate was placed on a microplate shaker at 300 rpm for 30 min at 4°C, and then incubated with monoclonal mouse anti-human RhoA primary antibody (# GL01A; 1:250; Cytoskeleton Inc.), followed by a polyclonal goat anti-mouse horseradish-conjugated secondary antibody (# GL02; 1:62.5; Cytoskeleton Inc.) on a shaker at 300 rpm at room temperature, for 45 min each. The plate was then incubated with the HRP detection reagent at 37°C for 15 min. After the addition of HRP stop buffer, absorbance was read at 490 nm using a microplate reader (Synergy H1, BioTek Instruments, Inc., Winooski, VT, USA).

### Western blotting

Western blot assay was used to examine the protein expression and/or activity of Piezo1, Cav-1, Src, ERK, FAK, and MMP-9. Cells grown on glass bottom dishes under described assay conditions were lysed using RIPA buffer (# R0278, Sigma) with added cocktail of protease and phosphatase inhibitors (MS-SAFE, Sigma). Protein concentration of cell lysates were determined using the Protein Assay Reagent (#23227, Thermo Fisher Scientific, Waltham, MA). Cell lysis buffer was combined in 4× SDS sample buffer and 2-mercaptoethanol and incubated at 95 °C for 5 min. After loading equal amount of protein per lane, SDS-PAGE was performed. The proteins were transferred onto 0.22 µm nitrocellulose membranes (# 66485, Pall Life Sciences) using Pierce G2 Fast Blotter (Thermo Fisher Scientific, Waltham, MA). Following transfer, the membranes were blocked using 5% nonfat milk in 1x TBST (Tris-buffered saline and 0.1% of Tween-20) for 1 h at RT with gentle agitation and incubated with the primary antibodies overnight at 4 °C under mild shaking condition. After washing three time with 1x TBST, membranes were incubated with goat anti-rabbit secondary antibody (DyLight 800, # SA5-10036, Thermo Fisher) or goat anti-mouse secondary antibody (DyLight 680, # 35518, Thermo Fisher) at RT for 1 h. Signal of immunoblots were detected using the Odyssey Infrared Imaging system (LI-COR, Lincoln, NE). For quantification, intensity of gel band was calculated after subtracting the background.

### Statistical analysis

Statistical analysis was done using two-tailed Student’s *t*-test. Statistical significance set to \**p* < 0.05 and \*\**p* < 0.01. All experiments were repeated at least three times and the data expressed as means ± s.e.m. (standard error of the mean).

## Results

### Compressive stress enhanced invasion of breast cancer cells dependent on Piezo1

Several methods exist to apply compressive stress to cells such as using a microfluidic platform^6,44^ or a hydrostatic pressure system^45,46^. However, a system that exerts a constant contact compressive force is more suitable to simulate uncontrolled growth-induced mechanical compression found in solid tumor. Thus we used a device with a piston of specific weight to apply a constant compressive force to breast cancer cells according to a previous report^14^. To test whether compressive stress enhances invasion of breast cancer cells, cells were grown on 2D membrane filter (8 μm pore) coated with Matrigel and covered with 1% of agarose gel, and then exposed to a constant weight (Figure 1a). The compressive stress levels used in this study were 200 Pa, 400 Pa, and 600 Pa, which were considered physiologically relevant as cells are reported to experience compressive stress at up to about 800 Pa in the core of the solid breast tumor^14,47^. As shown in Figure 1b and 1c, there were more MDA-MB-231 cells that invaded through the Matrigel-coated transwell filters when exposed to compressive stress of 200 Pa, 400 Pa, and 600 Pa as compared to their un-compressed counterparts (control). The results clearly show that compressive stress enhanced the breast cancer cell invasion in a pressure-dependent manner.

**Figure 1.**
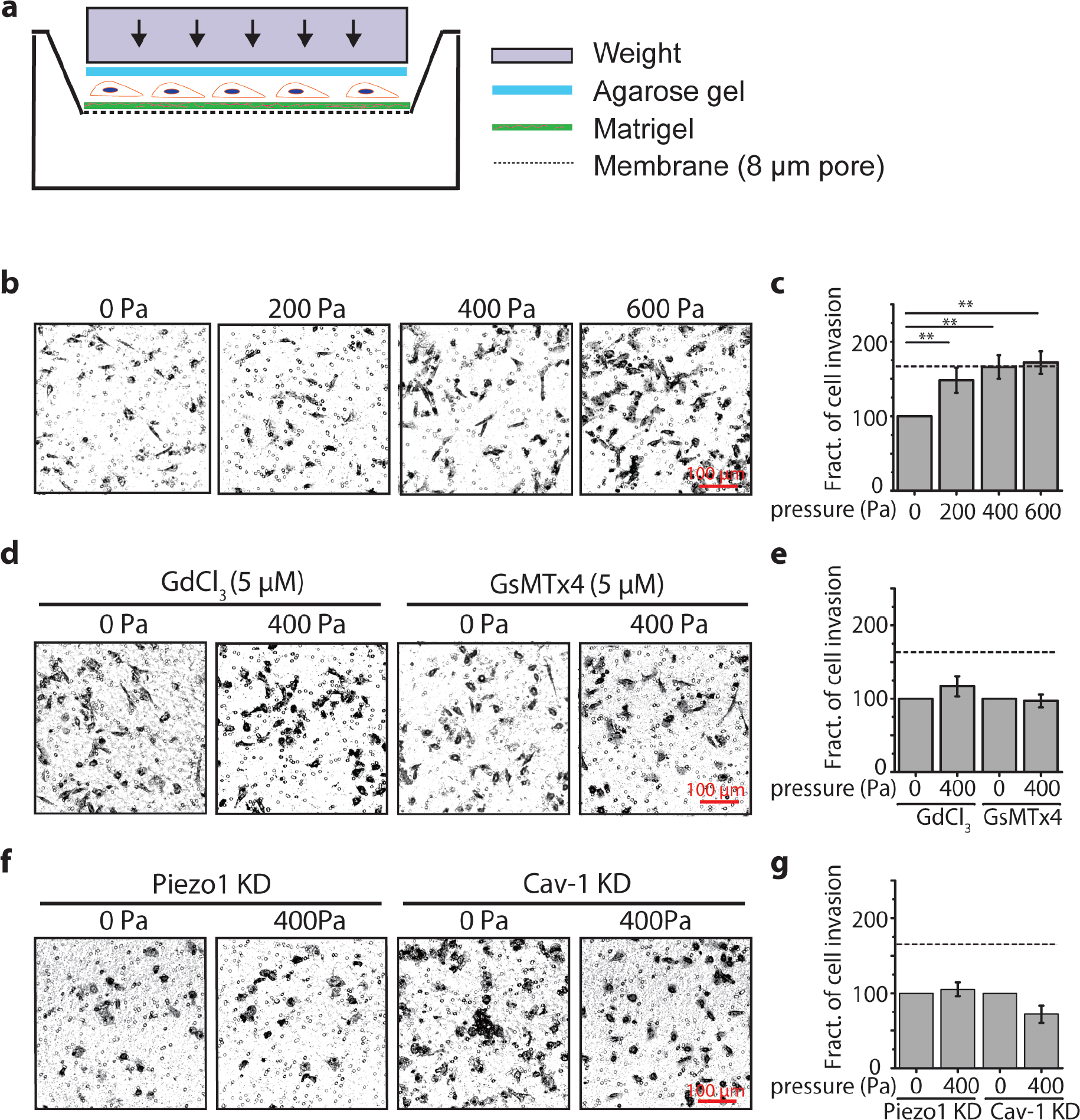
Compressive stress enhanced invasion of MDA-MB-231 cells depending on Piezo1. Cell invasion was measured with *in vitro* transwell invasion assay. **a** Schematic diagram of the compression experiment using a transwell setup. Cells grown on membrane filter (8 μm pore) coated with Matrigel for 6 h were covered with 1% of agarose gel and compressed with a specific weight. **b, c** Representative images of invaded cells stained with crystal violet under different compressive stress and quantification of percentage of invaded cells. **d**, **e** Representative images of invaded cells and quantification of percentage of invaded cells treated with Gadolinium chloride (Gd^3+^) or GsMTx4 under 400 Pa. **f**, **g** Representative images of invaded cells and quantification of percentage of invaded cells treated with siRNA for Piezo1 or Cav-1 under 400 Pa. Dashlines in **c**, **e**, and **g** indicate cell invasion under 400 Pa without any perturbations, n = 3.

We also found that the compressive stress had no such enhancement effect on migration of the normal breast cells (data not known), in consistency with others’ report^14^. To test whether this could be attributed to different Piezo1 expression level between cancer and normal cells, we examined the distribution and expression of Piezo1 in MDA-MB-231 cells (cancer) in comparison with that in MCF10A (normal) cells. We found that Piezo1 was expressed in MDA-MB-231 cells in the form of puncta structures and located not only on plasma membrane, but also over the intracellular space and nucleus (Figure S1a), which is consistent with data reported by Gudipaty *et al*.^48^. More importantly, Piezo1 expression in the cancer cells was ~3 fold higher compared to the normal cells (Figure S1b), supporting the possibility that Piezo1 may be the main compressive stress sensor in the cancer cells.

In order to test whether the compressive stress-enhanced cancer cell invasion was specifically mediated through SAC, and particularly Piezo1 channels, we first pretreated MDA-MB-231 cells with SAC inhibitors, Gd^3+^ or GsMTx4, followed by exposure of the cells to 400 Pa compressive stress. As shown in Figure 1d and 1e, pretreatment with Gd^3+^ or GsMTx4 either partially attenuated or completely abrogated the compressive stress-enhanced invasion of the breast cancer cells.

To further confirm the specificity of Piezo1 in mediating the compression-induced cancer cell invasion, we silenced the protein expression of Piezo1 in MDA-MB-231 cells by using siRNA prior to application of compressive stress to the cells. Western blot results confirmed that the efficiency of Piezo1 knockdown (KD) was ~70% (Figure S2a). When MDA-MB-231 cells pretreated with Piezo1 siRNA were exposed to 400 Pa compressive stress, the enhanced cell invasion in response to compression was completely abrogated (Figure 1f and 1g). This further supports Piezo1’s function in compressive stress-enhanced cancer cell invasion.

Previous work suggests that cholesterol content that directly influences membrane organization such as formation of a unique form of lipid rafts known as caveolae might regulate Piezo1 functions^49–53^. To test whether caveolae were involved in the compressive stress-enhanced cancer cell invasion, we silenced the expression of Cav-1 in MDA-MB-231 cells by about 60% (Cav-1 KD, Figure S2b) prior to subjecting the cells to compression. Consistent with our hypothesis, Cav-1 KD also abrogated the compressive stress-enhanced cancer cell invasion (Figure 1f and 1g).

### Compressive stress enhanced matrix degradation and invadopodia formation dependent on Piezo1

Cancer invasion requires the tumor cells not only to migrate but also to degrade their extracellular matrix in aid of the cells to move through dense barriers of their microenvironment. However, it is unknown whether compressive stress influences cancer cells’ capability for matrix degradation. To address this important issue, we examined the extent of matrix degradation and invadopodia formation of MDA-MB-231 cells seeded on FITC-conjugated gelatin-coated glass bottom dish followed by application of compressive stress. The fluorescence images showed dark puncta underneath the cells, corresponding to “holes” formed in the gelatin matrix due to degradation, and some of these holes were also colocalized with actin puncta formed on the cell membrane as described below (Figure 2a). Thus, we quantified the extent of matrix degradation underneath the MDA-MB-2312 cells, and the results showed that MDA-MB-231 cells exposed to compressive stress from 200 Pa to 600 Pa exhibited a significant and stress-dependent increase of gelatin matrix degradation as compared to their un-compressed counterparts (0 Pa) (Figure 2b and 2c). Similar as in the case of cell invasion through Matrigel-coated transwell filter, pretreatment of MDA-MB-231 cells with GsMTx4 to inhibit SAC or siRNA probe to silence Piezo1 expression completely abrogated the compressive stress-enhanced gelatin matrix degradation in the breast cancer cells, whereas silencing the expression of Cav-1 led to partial prevention of the compressive stress-enhanced gelatin matrix degradation at 400 Pa and 600 Pa, but not at 200 Pa (Figure 2c). These data indicate that compressive stress did enhance the degradation of ECM (in this case the gelatin) by the breast cancer cells in a Piezo1 dependent manner, and caveolae were involved in this process at least at high magnitude of compressive stress.

**Figure 2.**
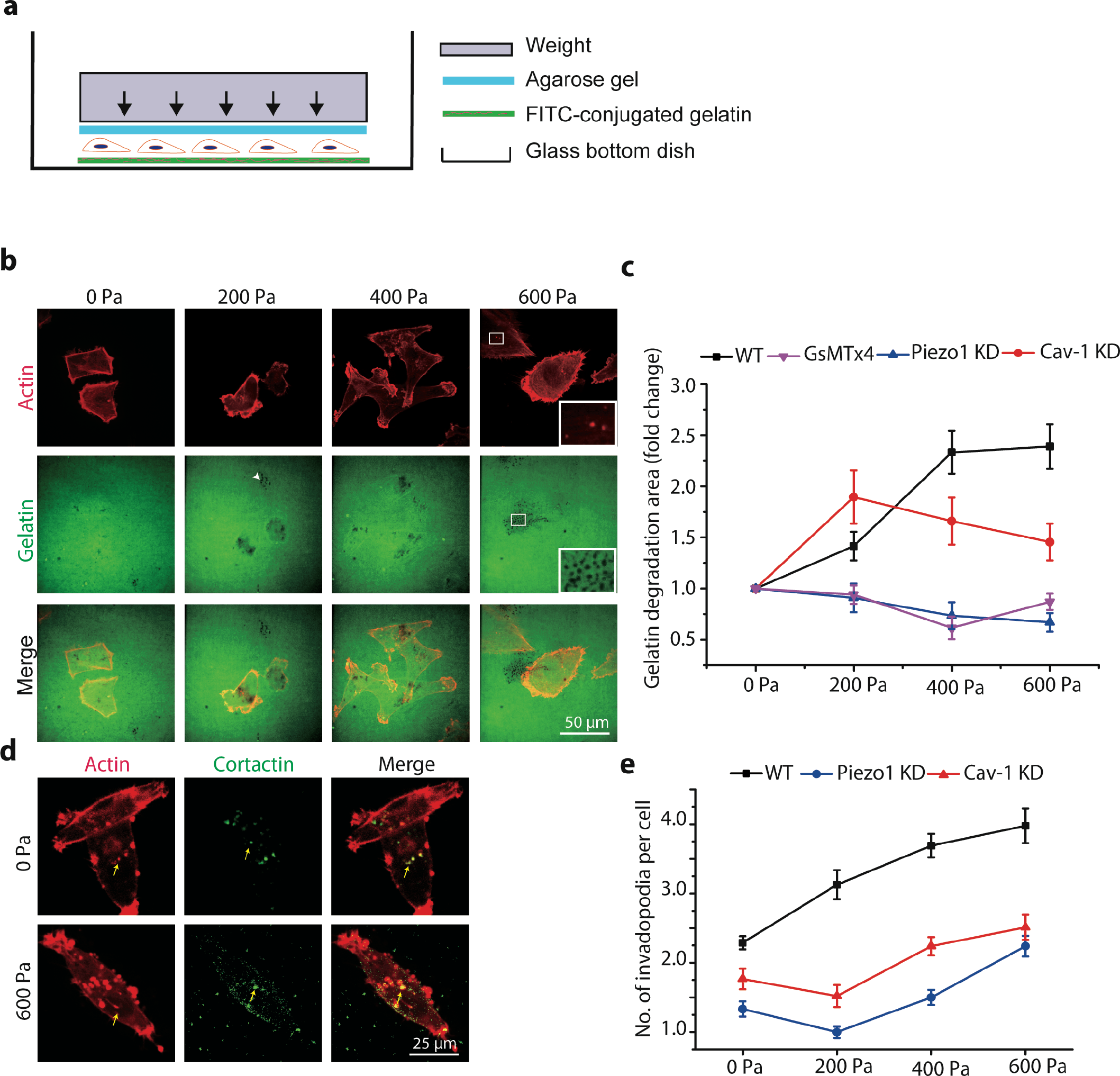
Compressive stress promoted matrix degradation and invadopodia formation in MDA-MB-231 cells. **a** Schematic diagram of the experiment. Cells grown on a glass-bottom dish coated with FITC-conjugated gelatin for 8 h were covered with 1% of agarose gel and compressed with a specific weight. **b** Representative images (red: actin, green: gelatin) of compression-promoted gelatin degradation at the ventral side of the cell. Gelatin degradation was visualized by confocal microscopy as disappearance of green fluorescence. The arrow in 200 Pa shows an example of the punctate gelatin-degraded region. Insets of 600 Pa images are magnified view of the boxed regions. **c** The fold change of gelatin degradation area under different treatment conditions as a function of compressive stress normalized to gelatin degradation area at 0 Pa. **d** Representative images (red: actin, green: cortactin) of invadopodia (yellow arrows) in MDA-MB-231 cells with or without 600 Pa compression. Control and compressive stress-treated MDA-MB-231 cells were cultured on gelatin-coated glass bottom dish for 8 h, then fixed and stained with phalloidin to visualize F-actin (red) and antibody for cortactin (green). **e** The number of invadopodia per cells under compression with or without Piezo1 KD or Cav-1 KD. n = 20-30 in **c** and **e**.

Cancer cells are known to use so-called invadopodia formed on the cell membrane as structural basis for initiation and promotion of ECM degradation^54^. These structures are dynamic actin protrusions which secret and localize enzymes required for ECM degradation and are controlled by both chemical and physical cues such as EGF, external mechanical force and matrix stiffness^55,56,57^. Thus, we examined whether compressive stress could also promote invadopodia formation in the cancer cells. We used immunofluorescence labeled cortactin, a marker for invadopodia^58^, to visualize and identify invadopodia as cortactin-positive actin puncta on the cell membrane (Figure 2d). The number of invadopodia per cell was manually counted and reported for MDA-MB-231 cells with or without pretreatment with siRNA probe to silence Piezo1 and Cav-1, respectively, and with or without exposed compressive stress. The results show that compressive stress increased the number of invadopodia per cell in MDA-MB-231 cells, which was significantly abrogated by silencing either Piezo1 or Cav-1 (Figure 2e). These results demonstrated that the breast cancer cells’ responded to compressive stress with increased number of membrane-based invadopodia that in turn promoted the ECM degradation, which essentially depended on the existence and function of both Piezo1 and caveolae. This finding is in agreement with the previously reported phenomenon of caveolae promoting invadopodia formation^33^, and support a working model that Piezo1 and caveolae mediate compressive stress-enhanced matrix degradation *via* invadopodia.

### Piezo1 mediated compressive stress-promoted formation of stress fibers and apical actin protrusions

Actin filaments serve as essential players in mechanotransduction and are highly dynamic in response to mechanical stress. Cells under compressive stress may reprogram their intracellular mechanical structures such as stress fibers to acquire appropriate mechanical properties. To observe the response of stress fibers to compressive stress, we generated a stable cell line expressing RFP-Lifeact and monitored the dynamics of actin filaments in MDA-MB-231 upon exposure to compressive stress. We observed frequently that compressive stress significantly promoted the formation of stress fibers at the cell cortex region in as early as 5 min in wild type cells (Figure 3a, top panel). In contrast, the compressive stress-enhanced formation of stress fiber was largely inhibited in the Piezo1 KD cells (Figure 3a, bottom panel). Interestingly, by using 3D scanning confocal imaging we discovered actin protrusions at the apical side of the MDA-MB-231 cells exposed to 600 Pa compressive stress (Figure 3b, upper panel), which were reduced in the Piezo1 KD cells exposed to the same amount of compressive stress (Figure 3b, bottom panel).

**Figure 3.**
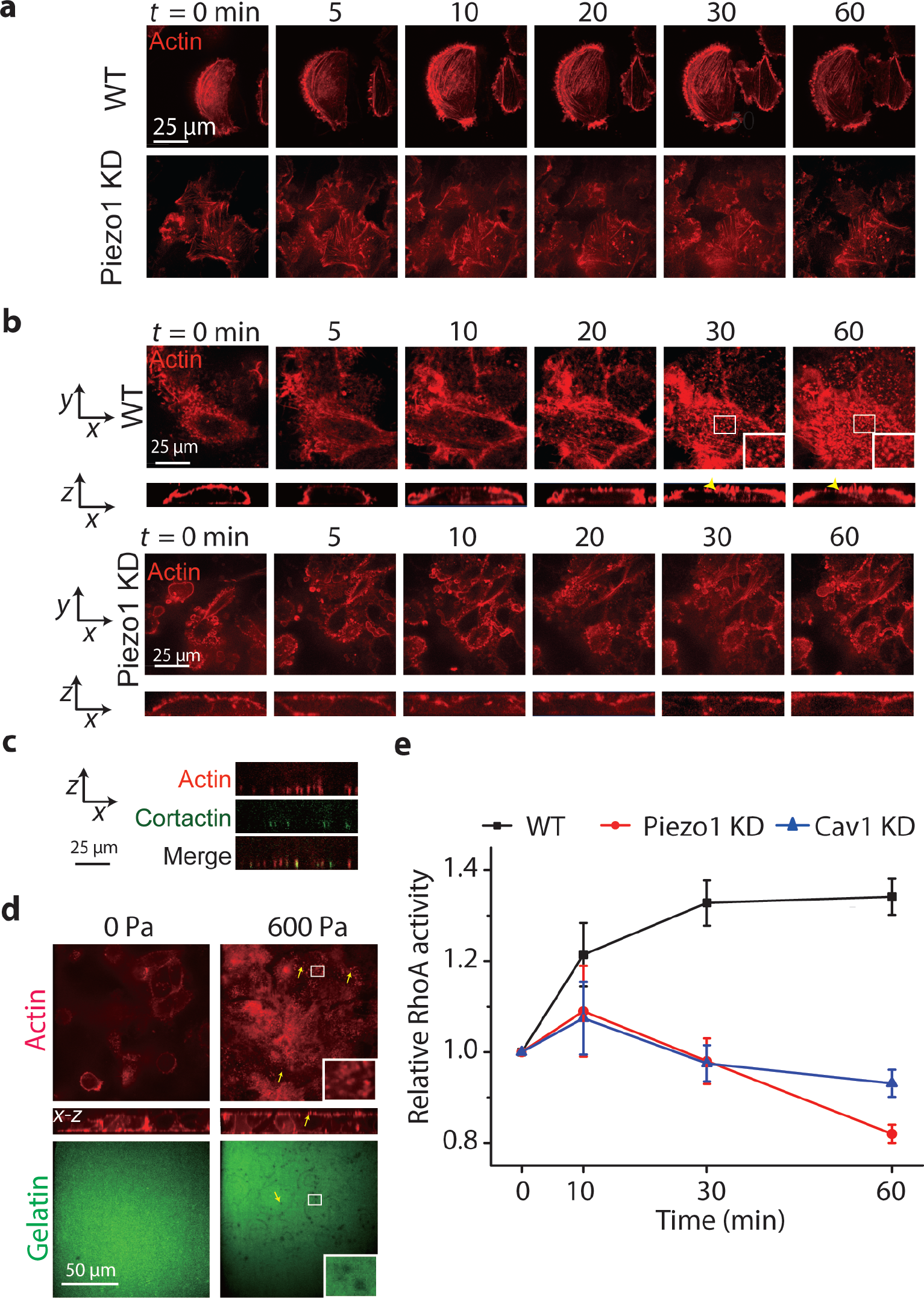
Piezo1 mediated compressive stress-induced formation of stress fiber and actin protrusions. **a** Representative images of stress fiber at the ventral side of MDA-MB-231 cells expressing Lifeact-RFP with or without Piezo KD under compression (600 Pa) for 60 min. **b** Representative images of actin protrusions at the apical side of MDA-MB-231 cells expressing Lifeact-RFP under compression (600 Pa) for 60 min (see magnified views and yellow arrow heads in the images at 30 and 60 min). **c** Representative images of cortactin with actin protrusion in MDA-MB-231 cells expressing Lifeact-RFP under compression (600 Pa) for 60 min. **d** Representative images of actin protrusions at the apical side of cells degraded gelatin. Insets of 600 Pa images are magnified view of the boxed regions. **e** G-LISA assayed relative RhoA activity of cells with different KD conditions being compressed at 600 Pa from 0 to 60 min, n = 3.

To our knowledge, such apical actin protrusions have not been reported before, but they appeared to be similar to the invadopodia found at the ventral side of the cells for mediating ECM degradation^56^. To verify whether or not these apical actin protrusions were invadopodia, we first examined the agarose gel that covered the cells during compression using immunofluorescence microscopy. We found that the apical actin protrusions that were stuck in the agarose gel colocalized with cortactin (Figure 3c). Then, we coated the agarose gel with FITC conjugated-gelatin and used it to cover the cells during compression. We found many dark puncta appeared in the gelatin at the apical side of the cells (as indicated by the yellow arrows in Figure 3d). These data together strongly suggest that the apical actin protrusions induced by compressive stress were similar, in both structure and function, to the invadopodia commonly found on the ventral side of the cells.

Since the formation of both stress fiber and invadopodia is determined by the dynamics of actin polymerization *via* Rho family GTPases, in particular RhoA signaling^59–61^, we used G-LISA to quantify the RhoA activity in MDA-MB-231 cells exposed to 600 Pa compressive stress for up to 60 min, and the cells were either wild type (WT), Piezo1 KD, or Cav-1 KD as described before. The results show that compressive stress activated RhoA activity in all cases within 10 min of compression, although the stress-activated RhoA activity was much greater in the WT cells compared to that in Piezo KD/Cav-1 KD cells. However, with continued compression, the RhoA activity in WT cells continued to increase until 30 min and then maintained at a plateau from 30 to 60 min, whereas that in the Piezo1 KD/Cav-1 KD cells turned to decrease persistently (Figure 3e). This suggests that the formation of stress fibers and invadopodia in the breast cancer cells in response to compressive stress was indeed mediated by Piezo1 and caveolae *via* RhoA signaling.

### Piezo1 mediated compressive stress-induced calcium signaling

Calcium signaling is essential in regulating cellular behaviors and can be activated by diverse mechanical stresses^62^. Therefore, it is important to determine whether calcium signaling was involved in the compressive stress-induced invasive phenotype of breast cancer cells in addition to the putative Piezo1 mechanism. We transiently transfected calcium biosensor G-GECO and performed live cell imaging during application of compressive stress to MDA-MB-231 cells. As shown in Figure 4a, calcium signaling in the cells, as indicated by the fluorescence intensity of G-GECO, was activated instantaneously upon exposure to compressive stress (**Video 1**). At about 1 min of compression, the magnitude of activation (the mean value of fluorescence intensity of G-GECO) peaked and the peak magnitude also increased from ~1.5 to ~4 fold as the compressive stress increased from 200 Pa to 600 Pa (Figure 4b).

**Figure 4.**
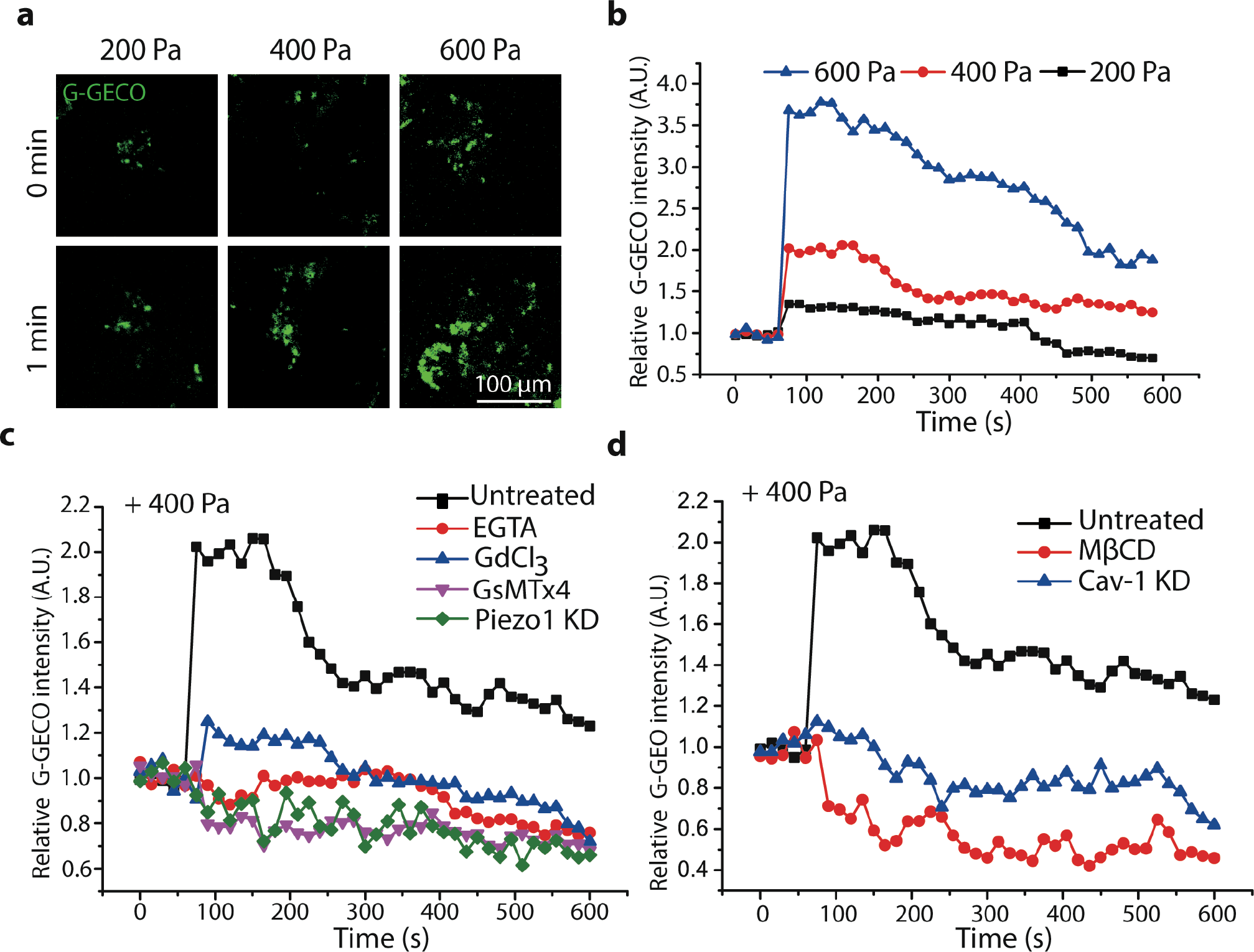
Compressive stress induced calcium signaling in MDA-MB-231 cells. **a** Representative images of intracellular [Ca^2+^] in cells stably expressing G-GECO before (0 min) and after (1 min) exposure to compressive stress at 200, 400, 600 Pa, respectively. **b-d** Time-courses of changing relative mean fluorescence intensity of G-GECO (normalized to time 0) in MDA-MB-231 cells in response to compressive stress at 200, 400, and 600 Pa, respectively; in MDA-MB-231 cells pretreated with or without EGTA, Gd^3+^, GsMTx4, Piezo1 KD, MβCD and Cav-1 KD in response to 400 Pa compressive stress. Each experiment assayed 10-20 cells and repeated three times.

Moreover, we treated the cells with 2 mM EGTA for 15 min to deplete the extracellular calcium content before application of compressive stress (400 Pa), which completely eliminated the compressive stress-induced calcium signaling, suggesting the signaling was mainly due to influx of extracellular calcium rather than intracellular calcium activation (Figure 4c). Furthermore, the calcium influx induced by compressive stress (400 Pa) was also abrogated when the cells were pretreated with Gd^3+^ or GsMTx4 to block SAC, or siRNA probe to silence Piezo1 expression (Figure 4c). Finally, when the cells were pretreated with either 5 mM MβCD that dramatically reduced the number of caveolae in the cells (Figure S3), or siRNA probe to silence Cav-1 expression in the cells (Cav1 KD), the compressive stress-induced calcium influx was blocked (Figure 4d). Together, these observations support the finding that Piezo1 activity was dependent on intact caveolae and Piezo1 mediated the cellular response to compressive stress *via* calcium influx.

### Piezo1 mediated compressive stress-enhanced Src/ERK activation

During invadopodia formation and maturation to degrade matrix, several signaling pathways are involved including Src/ERK pathways^63^ which promote the formation of invadopodia^64^, and FAK pathway which promotes matrix metalloproteinase (MMP) synthesis and secretion^65^. Many investigations have shown that MMP-9 are highly expressed in breast cancer cells which contribute to the degradation of basal membrane that strongly correlated with breast cancer progression^66,67^. Thus, MMP-9 might be associated with the change of invasion and metastasis under compression. To test whether these signaling pathways are activated by compressive stress to regulate formation and function of invadopodia, we quantified the phosphorylation of ERK, Src, FAK and the expression level of MMP-9 in MDA-MB-231 cells following compression. We found that compressive stress significantly activated ERK, Src and FAK and promoted the expression of MMP-9 (Figure 5a). Piezo1 KD effectively abolished the compressive stress-promoted signaling of most pathways, except for ERK, and diminished MMP-9 expression (Figure 5b). The elevated ERK under compressive stress in Piezo1 KD may be due to pathways that do not depend on calcium signaling.

**Figure 5.**
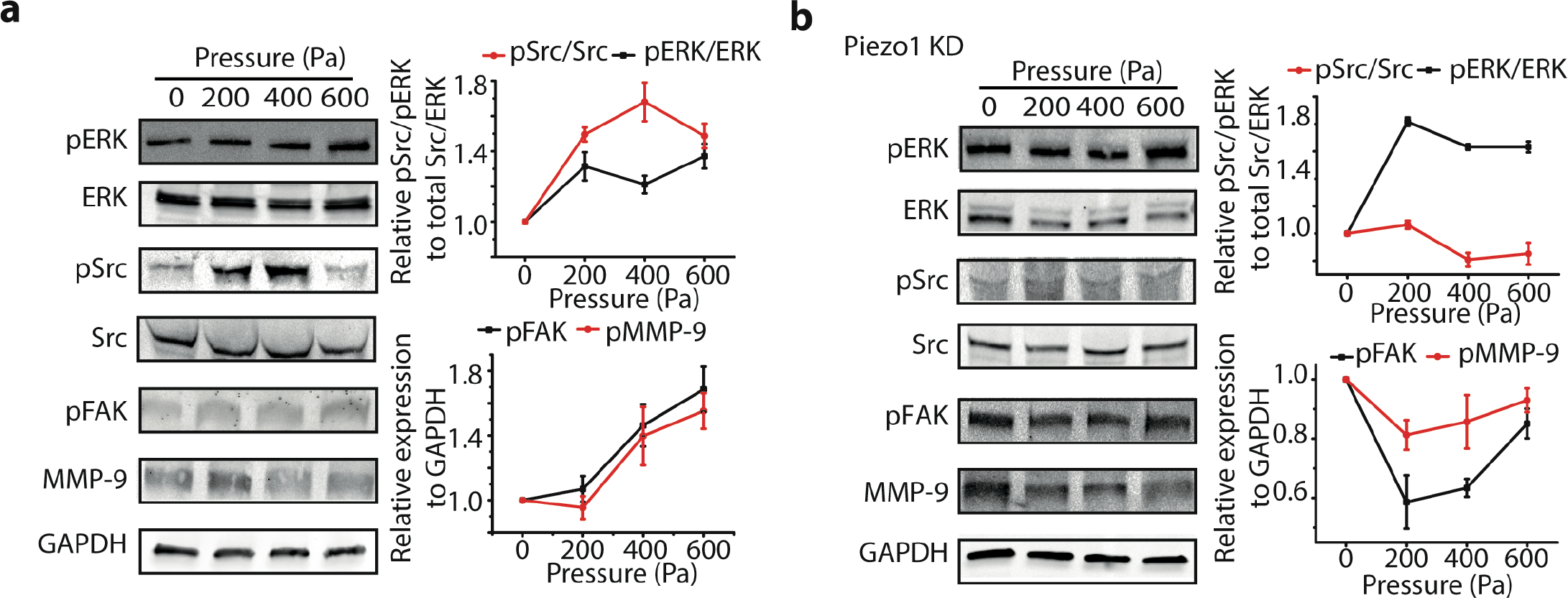
Compression enhanced the activity of ERK, Src, FAK and the expression of MMP-9. **a** The phosphorylation and expression of ERK, Src, FAK and MMP-9 in MDA-MB-231 cells in the absence or presence of compressive stress at 200, 400, 600 Pa. **b** The phosphorylation and expression of ERK, Src, FAK and MMP-9 in MDA-MB-231 cells pretreated with siRNA for Piezo1 in the absence or presence of compressive stress at 200, 400, 600 Pa. Relative phosphorylation or expression levels were obtained by normalizing to GAPDH expression and 0 Pa value, n = 3.

### Caveolae regulated the protein expression and distribution of Piezo1

The dependence of invasive phenotypes on intact caveolae led us to speculate whether Piezo1 might be functionally associated with caveolae. In both wild type MDA-MB-231 cells (Figure 6a, upper panel) and the same type of cells transiently transfected with Cav-1 EGFP (Figure 6a, bottom panel), we found that both Piezo1 and caveolae formed puncta structures and many of them were colocalized. Interestingly, Cav-1 EGFP^+^ exhibited larger puncta structures compared with Cav-1 EGFP^-^ cells, suggesting that Cav-1 may regulate expression level of Piezo1. To test this hypothesis, we quantified Piezo1 protein expression level in wild type, Cav-1 EGFP transiently transfected, and Piezo1 KD MDA-MB-231 cells. Consistent with the idea that Cav-1 regulates Piezo1 expression, Cav-1 EGFP expression increased the Piezo1 expression while Cav-1 KD decreased the Piezo1 protein expression (Figure 6b). The functional relationship between Piezo1 and caveolae was further examined by addition of 5 mM of MβCD which disrupts caveolae in the cells. Consequently, the fluorescence intensity of Piezo1 was reduced in the cytosol but increased in the nucleus at the 5 min time point, suggesting Piezo1 translocation to the nucleus when caveolae were disrupted (Figure 6c). Thus, caveolae may regulate the organization and distribution of Piezo1 in breast cancer cells.

**Figure 6.**
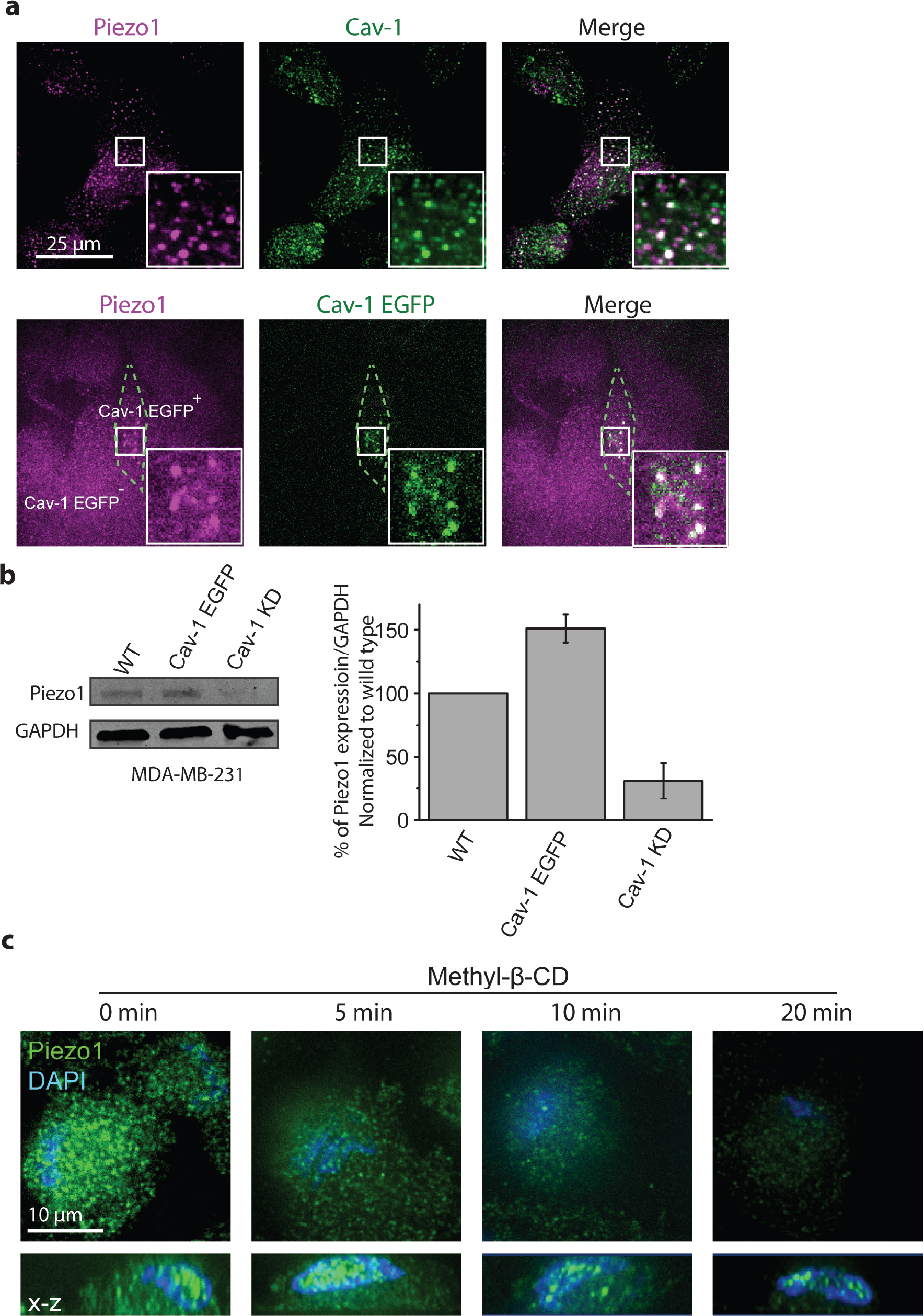
The expression and distribution of Piezo1 in MDA-MB-231 cells were regulated by caveolae. **a** Representative fluorescence images of Piezo1 (magenta) and caveolae (green) colocalization in endogenous (top panel) and Cav-1 EGFP expressing (bottom panel) MDA-MB-231 cells. Cav-1 EGFP positive and negative cells are present and labeled accordingly. Insets in both conditions show magnified view of the boxed regions. **b** Western blot images and quantification of Piezo1 expression in WT, Cav-1 EGFP expressing, and Cav-1 KD MDA-MB-231 cells (n = 3). **c**, Representative fluorescence images of Piezo1 (green) and nucleus (blue) after cells were treated with MβCD for 5 min, 10 min and 20 min (upper panel: *x*-*y* view, lower panel: *x*-*z* view).

## Discussion

Compressive stress is a fundamental feature of cancer microenvironment^68^. Many reports have shown that compressive stress in cancer affects the efficiency of tumor treatment not only by influencing the distribution of drugs but also by affecting cancer cell behaviors^13^. Giving the importance of compressive stress in cancer development and treatment, substantial efforts are needed to understand how compressive stress affect the behaviors of cancer cells and its underlying mechanisms. In the current study, we first observed that in breast cancer cells, simulating uncontrolled growth-induced compressive stress enhanced cell invasion, matrix degradation, and invadopodia and stress fiber formation. Additionally, we identified that Piezo1 channels mediated these processes and this depends on Piezo1’s functional association with caveolae. These findings provide the first demonstration that compressive stress due to uncontrolled growth enhances matrix degradation by breast cancer cells and Piezo1 channels are an essential mechanosensor and transducer for such stress in breast cancer.

Invasion of cells through layers of ECM is a critical activity during cancer metastasis. Uncontrolled growth-induced compressive stress triggers invasive phenotype by forming leading cells and enhancing cell migration which in turn facilitates cancer progression^14^. It is also known that the capability of forming invadopodia is directly correlated with their invasive potential in breast tumor cells^69^. It is, however, unknown whether such stress in solid tumor would affect the capability of cancer cells to induce ECM degradation. Consistent with enhanced invasion under compressive stress, we found compressive stress enhanced matrix degradation *via* the activation of signaling pathways leading to invadopodia formation. Thus, it is plausible that compressive stress that increases with proliferation of cancer cells in solid tumor might initiate the invasion by enhancing the capability of matrix degradation *via* invadopodia formation. If that is the case *in vivo*, compressive stress might promote cancer cells to ‘dig more holes’ in the basement membrane which provide a way for their metastasis.

While it is well known that compressive stress affects cancer progression, how cancer cells sense and respond to such stress is not completely understood. Cell membrane exhibits fluid-like mechanical properties and has a well-defined membrane tension. Many reports have shown that mechanical stress increased membrane tension and regulate cell function^70^. Under compressive stress, cell membrane will be stretched which, in turn, increase the membrane tension. Piezo channels are the most notable family of SAC in mammalian cells which are gated by membrane tension^71^. It has been found in breast cancer cells that Piezo1 channels play essential roles in diverse pathological processes including cancer cell migration and the Piezo1 mRNA expression level is highly correlated with the survival time of breast cancer patients^29^. Our study also confirmed that Piezo1 channels are highly expressed in breast cancer cells compared to normal breast cells, and is consistent with the data that breast cancer cells but not normal breast cells respond to compressive stress regarding cell migration^14^.

Emerging evidence indicate that caveolae, subdomains of the plasma membrane that contain high concentration of cholesterol and sphingolipid, harbor and modulate receptors and ion channels. TRPV1 channels expression and distribution on the plasma membrane can be disrupted by the removal of caveolae *via* cholesterol depletion by MβCD^72^. Similar to TRPV1, we found Piezo1 behaves similarly by shifting its localization to the nucleus upon cholesterol depletion. Interestingly, in stretch-triggered mitosis, Piezo1 was also observed to localize to the nuclear envelope^48^. Thus, it may be a general strategy where a functional relationship between caveolae and Piezo1 regulates force sensing and our data supports the model where caveolae might be the “mechanical force foci” which concentrates Piezo1 to facilitate force sensing and transduction in mammalian cells.

The mechanisms how Piezo1 channels are gated by mechanical stress is still unclear. Piezo1 channels appear to sense tension in the bilayer and are gated according to “Force-from-lipid” principle, an evolutionally conserved gating mechanism^22^. According to this paradigm, the activity and sensitivity of Piezo1 channels can be regulated by lipid membrane and its physical properties. The physical properties of lipid membrane such as thickness, stiffness, and pressure profile found within caveolae may be different from the surrounding membrane. For instance, membrane stiffness is reduced by cholesterol depletion^50^, and disruption of caveolae integrity by cholesterol depletion has been demonstrated to change membrane stiffness and result in suppression of some epithelial sodium channels (ENaC) and TRP channels^52,73,74^. In this context, it is plausible that cholesterol-enriched caveolae might affect sensitivity of Piezo1 *via* controlling the membrane pressure profile. Consistent with membrane organization/composition playing a key role, recent studies also show that phosphoinositide depletion inhibits the activity of Piezo channels^49^. Stomatin-like protein-3 (STOML3) has been reported to tune the sensitivity of Piezo1 channels by controlling the membrane mechanical properties through recruiting cholesterol^50^. Taken together, it is likely that Piezo1 is located in cholesterol-rich caveolae microdomains and caveolae integrity regulates Piezo1 function.

The close relationship between Piezo1 and caveolae is perhaps not entirely surprising. Piezo1 channels are known to sense changes in tension rather than absolute tension in the cell membrane^20^. Caveolae are also recognized as being important in sensing external stimuli and have been shown to serve as membrane reservoirs to compensate mechanical perturbations such as osmotic stress and stretch^34,35^. Therefore, the flask-like invagination of caveolae may provide a situation in which the change of membrane tension is much larger when cells are exposed to compression.

In addition to membrane structures, cells upon mechanical stress can also reorganize their cytoskeletal structures to adapt new mechanical microenvironment. Among them, the stress fibers are essential mechanical structures which control various cellular behaviors. Reports have shown that mechanical tension induces the assembly of stress fibers^75,76^. In this study, we found that cells under compressive stress quickly assembled new stress fibers inside cells as early as in 5 min. This may be to increase their cellular mechanical strength to balance compressive stress. There is strong evidence that the assembly and organization of mechanically induced stress fibers are mediated by RhoA^60^, and mechanical stretch can activate RhoA within 4 min^77^, which are all consistent with what we have shown.

Interestingly, we also found that compressive stress induced actin protrusions at the apical side of cells which has never been reported previously. Our results support that these are invadopodia, as they contain cortactin and possess the ability to degrade gelatin. Consistent with this, these apical actin protrusions were absent in Piezo1 KD cells in which the elevated Src activity was also abolished. Our results also demonstrate the regulatory roles of several key pathways of mechanotransduction in the compressive stress-induced cancer cell invasive phenotype including MMPs, FAK/ERK and calcium, which are consistent with that mechanical force stimulates expression of MMPs, as well as growth factors, often utilizes FAK or ERK signaling pathway, which are activated by calcium signaling^78,79,80^. Our study thus provides a comprehensive understanding from disparate systems in the context of compression-induced breast cancer cell invasion (Figure 7).

**Figure 7.**
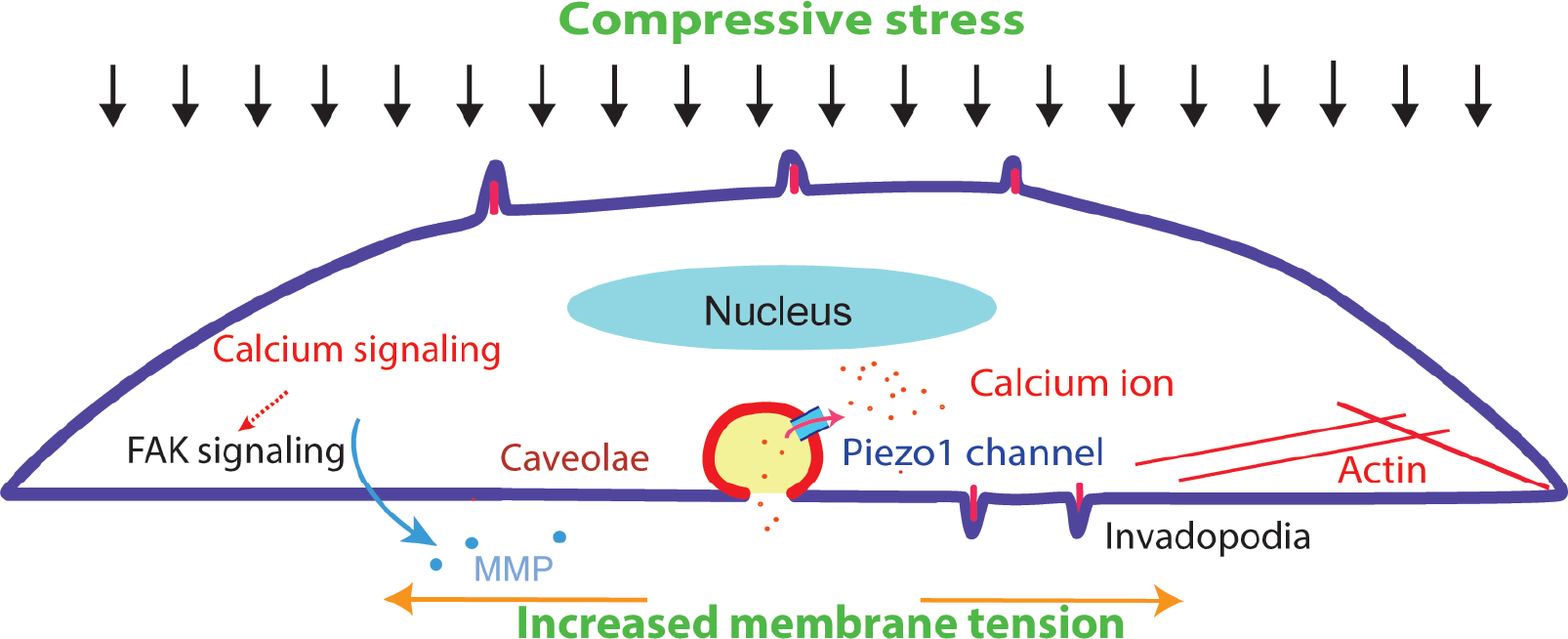
Model of compressive stress-promoted invasive phenotype of MDA-MB-231 cells and associated signaling pathways.

Our work may have relevance to human tumors *in vivo*. As solid tumor experiences high compressive stress due to uncontrolled proliferation and confinement by the stiff extracellular matrix environment, this microenvironment facilitates compression-enhanced cell invasion. The identification of Piezo1’s crucial role in this process provides the first demonstration of the dependence of Piezo1 channels on the response of breast cancer cells to physiological compressive stress. The functional dependence of Piezo1 on caveolae further highlights the importance of membrane organization and composition on force-gated ion channels. Both of these findings underscore the cardinal role that Piezo1 channels play in regulating cell invasion and may inspire further development targeting Piezo1 as a potential cancer therapeutic target.

## Acknowledgements

We thank Takanari Inoue (Johns Hopkins University) for providing the G-GECO plasmid. The technical assistance from Shue Wang and Maxwell DeNies is gratefully acknowledged. Funding for the work was provided by the Key Program of NSF of China (No. 11532003) to L.D. M.L. is supported by Jiangsu Oversea Visiting Scholar Program for University Prominent Young and Middle-aged Teachers and Presidents. A.P.L. is supported by NSF-MCB 1561794.

## Author contributions statement

M.L conceived, designed, and performed the experiments, analyzed the data, prepared figures, and wrote the manuscript. K.H., Z.T., and L.D. revised the manuscript. A.P.L. conceived the study and wrote the manuscript. All authors reviewed the manuscript.

