## Supplemental Figures for "Compressive Stress Enhances Invasive Phenotype of Cancer Cells via Piezo1 Activation"

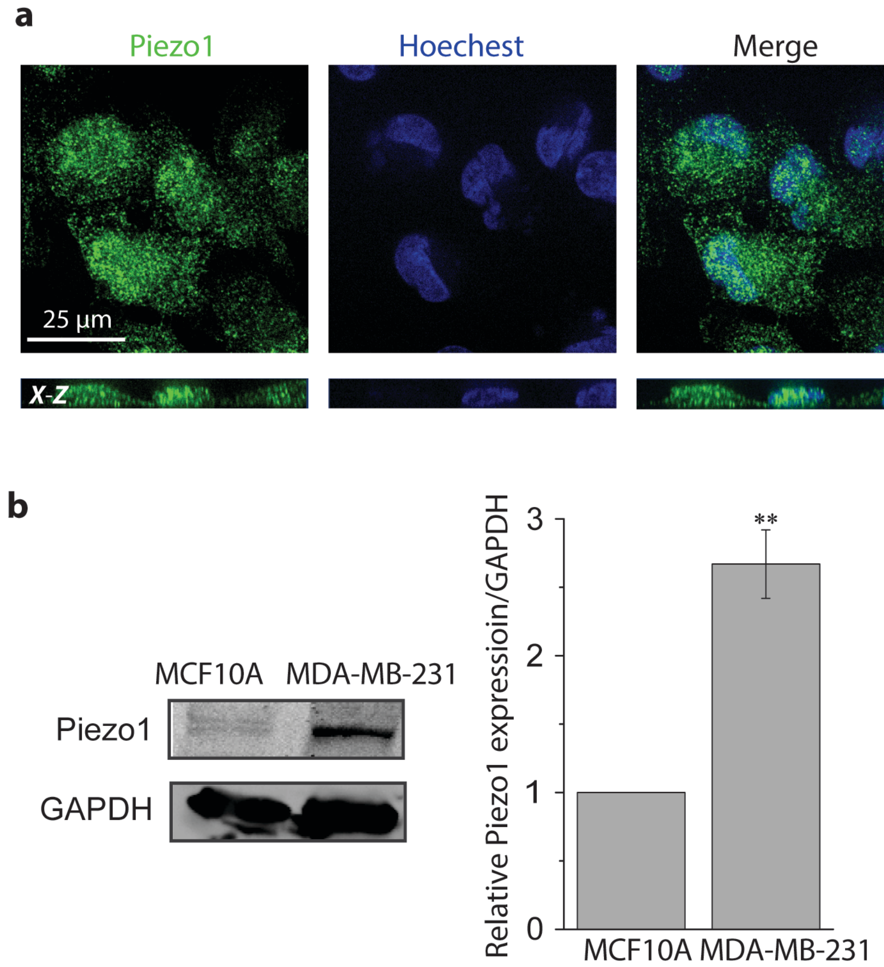


**Figure S1** Piezo1 distribution and expression in breast cancer or normal breast cells. **a** Representative confocal microscopic images of Piezo1 in MDA-MB-231 cells (100X), Piezo1 (green) and nucleus (blue) were detected with antibodies and Hoechest 33342, respectively, showing puncta structures of Piezo1 localized in plasma membrane, cytosol, and nucleus. **b** Western blot images and quantification of protein expression of Piezo1 in MCF10A (normal breast) and MDA-MB-231 (cancer breast) cells (** $p < 0.01$, n=3).

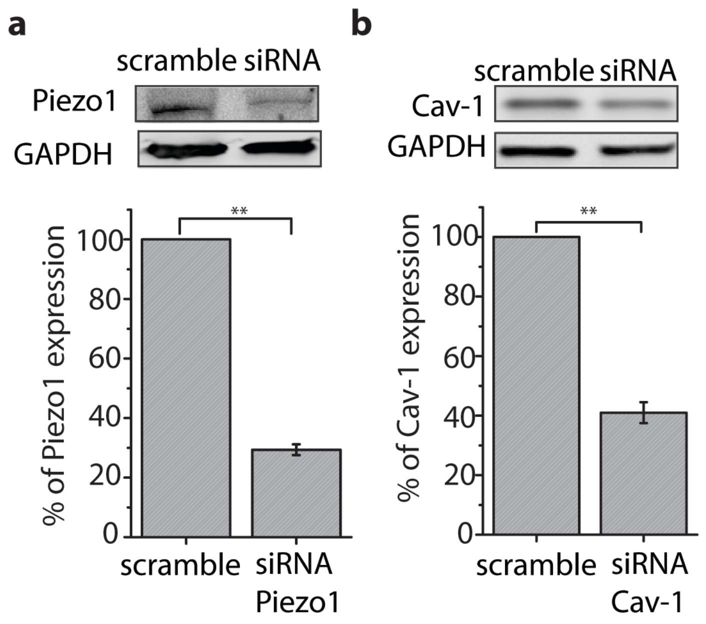


**Figure S2** Efficiency of siRNA KD for Piezo1 or Cav-1. The cells were transfected with scramble control or siRNA probes for Piezo1 (**a**) or Cav-1 (**b**) for 48 h and Piezo1 or Cav-1 expression was determined by Western blot. The bar graphs show the percentage of Piezo1 (**a**) or Cav-1 (**b**) in scramble and siRNA groups (** $p$ <0.01, n=3).

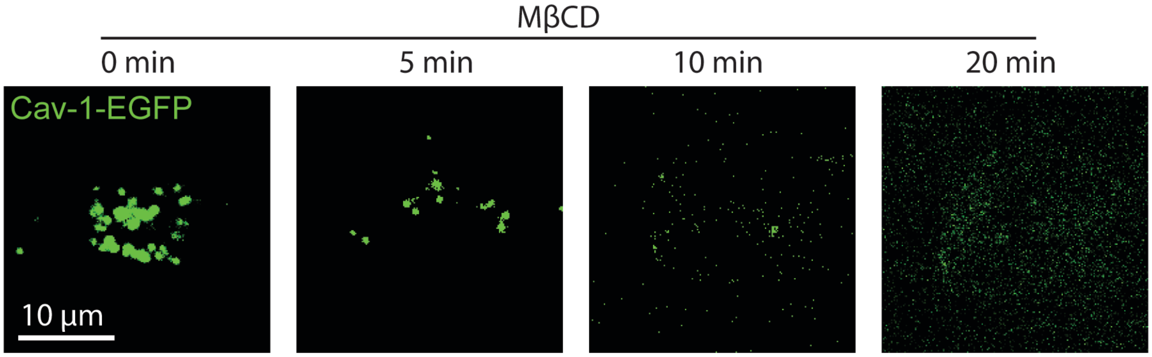


**Figure S3** Effect of methyl-beta-cyclodextrin (MβCD) on removing caveolae from the plasma membrane. Cells expressing Cav-1-EGFP were treated with 5 mM of MβCD for 5 min, 10 min, and 20 min at 37 °C and imaged with confocal microscope (100X). The representative images show that MβCD dramatically removed caveolae from the plasma membrane within 20 min of the drug administration.
